# Natural products from food sources can alter the spread of AMR plasmids in *Enterobacterales*

**DOI:** 10.1101/2024.02.26.582065

**Authors:** Ilyas Alav, Parisa Pordelkhaki, Judith Rodriguez-Navarro, Onalenna Neo, Celia Kessler, Ruth Jesujobalayemi Awodipe, Poppea Cliffe, Huba Marton, Simon Gibbons, Michelle M.C. Buckner

## Abstract

Antimicrobial resistance (AMR) poses a significant threat to global public health. Notably, resistance to carbapenem and extended-spectrum β-lactam antibiotics in Gram-negative bacteria is a major impediment for the treatment of infections. Genes responsible for resistance to these antibiotics are frequently carried on plasmids, which can transfer between bacteria. Therefore, exploring strategies to prevent this transfer and/or the prevalence of AMR plasmids is timely and pertinent. Here, we show that certain natural product extracts and associated pure compounds can reduce the transmission of AMR plasmids into new bacterial hosts. Using our established high-throughput fluorescence-based screen we found that the natural products were more active in reducing transmission of the IncK plasmid pCT in *Escherichia coli* ST131, compared to *Klebsiella pneumoniae* Ecl8 carrying the IncFII plasmid pKpQIL. Furthermore, we found that the natural product rottlerin was more active in *K. pneumoniae* than in *E. coli*. Importantly, rottlerin was also associated with a reduced number of transconjugant bacteria in a clinical *K. pneumoniae* isolate harbouring a *bla*_NDM-1_ plasmid. Together, these results demonstrate the potential of natural products as promising anti-plasmid agents.

## Introduction

Antimicrobial resistance (AMR) is a growing global problem, with resistant bacteria causing increasing numbers of difficult-to-treat infections leading to increased morbidity and mortality. Of particular concern are extended-spectrum beta-lactamase (ESBL) and carbapenemase-producing Enterobacteriaceae (1). One of the factors that has resulted in the widespread dissemination of ESBLs and carbapenemases amongst these organisms are mobile genetic elements, such as plasmids. Plasmids are self-replicating pieces of DNA which can carry a variety of accessory genes, including multiple AMR and/or virulence genes (2, 3). Conjugative plasmids encode all the necessary machinery to mediate their transmission from one bacterial cell to another through a process called conjugation (4). Conjugative transfer of a plasmid into a new host has the potential to turn a drug-susceptible strain into a multidrug-resistant strain and in worst-case scenarios, also a hypervirulent strain (5). Therefore, research into these mobile genetic elements is of considerable importance.

Some approaches have focused on different methods to remove plasmids from bacterial hosts (curing agents), and others have focused on preventing plasmid transfer (conjugation inhibitors); broadly speaking such anti-plasmid approaches are gaining interest (6, 7). The search for and use of anti-plasmid compounds capable of curing plasmids or inhibiting conjugative plasmid transfer is ongoing (8-11). Natural products have historically played a significant role in drug discovery owing to the extensive structural diversity and complexity of chemical compounds (12). Certain food products and their bioactive constituents possess diverse physiological effects. For example, ginger has been reported to have protective effects on gastrointestinal, nervous, and cardiovascular systems (13), displays antimicrobial effects (14), and has been associated with improved outcomes in fatty liver diseases (15). Similarly, black pepper and turmeric extracts have been reported to have diverse physiological effects *in vitro* and *in vivo*, including anti-tumorigenic, anti-diarrhoeal, antioxidant, and antimicrobial effects (16, 17). The kamala tree (*Mallotus philippensis*) and its fruit have been traditionally used to treat parasitic infections and reported to have antimicrobial, antioxidant, and anti-inflammatory properties (18). Therefore, the wealth of diverse phytochemicals in natural products could offer compounds with anti-plasmid activity.

Here, we performed a screen for natural products with anti-plasmid activity (with either plasmid curing or conjugation inhibitor activity) by measuring the effects of bioactive plant extracts and bioactive compounds from black pepper, ginger, turmeric, cashew nuts and kamala extracts on plasmid transfer in *Escherichia coli* and *Klebsiella pneumoniae*.

## Results

### Extracts from natural products can reduce plasmid transmission

Given the highly bioactive properties of certain food products, it was decided to test the activity of extracts from black pepper, ginger, turmeric, and kamala using a previously developed high-throughput screening assay (8) of pCT and pKpQIL transmission in *E. coli* and *K. pneumoniae*, respectively (Fig. 1). For pCT transmission in *E. coli,* generation of transconjugants was reduced by all extracts compared to the DMSO control: black pepper 14.0% of the DMSO control (*p*=0.0007), ginger 20.9% (*p*=0.0019), turmeric 25.8% (*p*=0.0046), and kamala 28.9% (*p*=0.0090) (Fig. 1A). However, pKpQIL transmission in *K. pneumoniae* was generally less affected by the natural products. Black pepper reduced transconjugant production to 37.4% of the DMSO control (*p*=0.0352), turmeric reduced to 20.6% (*p*=0.0024), and kamala reduced to 31.6% (*p*=0.0099). However, ginger extract had no significant effect (112%, *p*=0.7227) (Fig. 1b). Comparing the impact between pCT transmission in *E. coli* and pKpQIL transmission in *K. pneumoniae*, turmeric and kamala extracts had relatively comparable effects, while ginger and black pepper were less active in *K. pneumoniae* than in *E. coli*.

**Figure 1.**
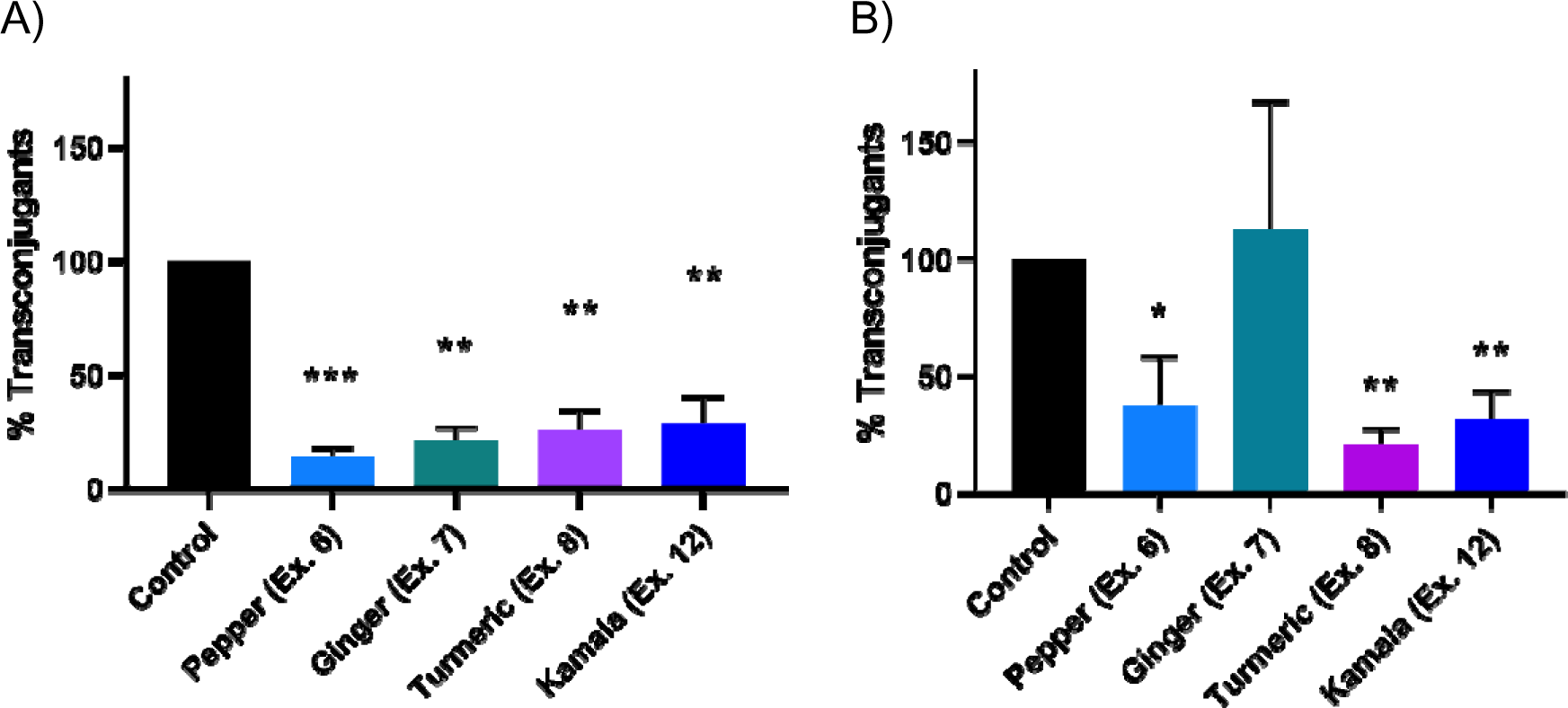
Effect of natural product extracts at 0.25 mg/mL on transmission of **A)** pCT in *Escherichia coli* and **B)** pKpQIL in *Klebsiella pneumoniae*. Data shown are the mean ± standard deviation from three independent experiments, each carried out with four biological replicates. The percentage of transconjugants following incubation with extracts was compared to DMSO control using a paired *t*-test. Significantly different results are indicated with * (*p* ≤ 0.05), ** (*p* ≤ 0.005), or *** (*p* ≤ 0.001).

### Pure compounds from natural product extracts have a moderate effect on plasmid transfer

Based on the literature, some pure compounds with known bioactive effects found in the food product extracts, or compounds with anticipated activity were tested using the high-throughput conjugation assay (Fig. 2). As with the whole extracts, pCT transmission in *E. coli* was more susceptible to inhibition than pKpQIL transmission in *K. pneumoniae*. All compounds were tested at 100 µg/mL. The data showed that the percentage of pCT transconjugants produced was significantly reduced to 43.7% (*p*=0.0039) for 6-gingerol, 21.4% (*p*=0.0069) for capsaicin, 106% (*p*=0.7055) for anacardic acid, and 89.1% (*p*=0.0168) for rottlerin, compared to DMSO control (Fig. 2A). While in *K. pneumoniae*, rottlerin was the only compound that significantly reduced production of pKpQIL transconjugants to 49.7% (*p*=0.0291) compared to the DMSO control (Fig. 2B). Surprisingly, capsaicin and anacardic acid showed an increase in pKpQIL transconjugant production (140%, *p*=0.0478 and 131% *p*=0.0419, respectively) compared to DMSO control. 6-gingerol had no significant impact (98.3%, *p*=0.8614) on the number of pKpQIL transconjugants compared to the control (Fig. 2B).

**Figure 2.**
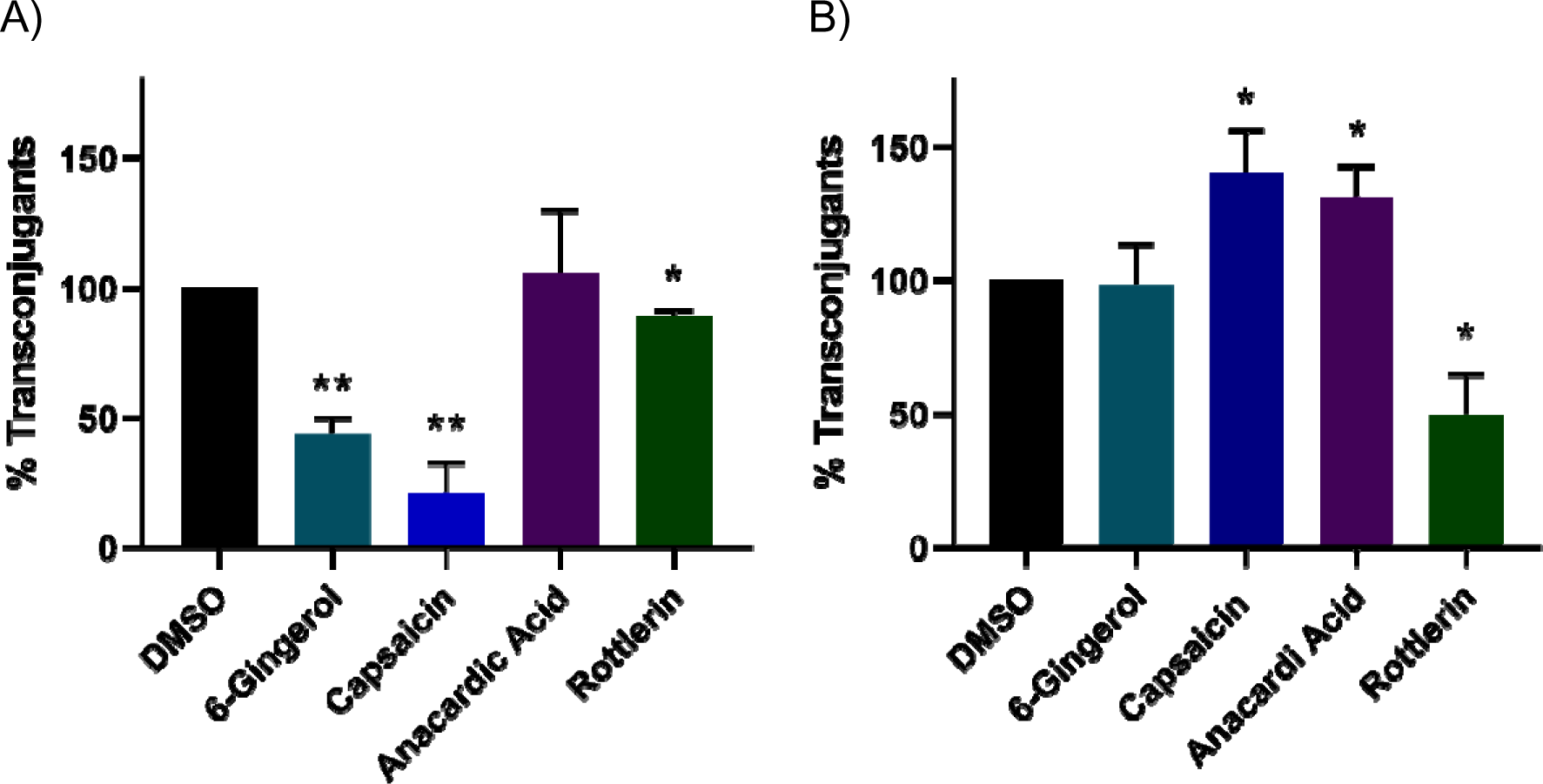
Impact of pure compounds found in the natural product extracts at 100 µg/mL on the transmission of **A)** pCT in *Escherichia coli* and **B)** pKpQIL in *Klebsiella pneumoniae*. Data shown are the mean ± standard deviation from three independent experiments, each carried out with four biological replicates. The percentage of transconjugants following incubation with compounds was compared to a DMSO control using a paired *t*-test. Significantly different results are indicated with * (*p* ≤ 0.05) or ** (*p* ≤ 0.005).

To look in more detail at the impacts of the two compounds with the greatest activity in *E. coli*/pCT transmission, dose-response curves were performed with concentrations of 1-256 µg/mL of 6-gingerol and capsaicin. These experiments showed a dose-dependent reduction in production of pCT transconjugants at concentrations of 6-gingerol above 64 µg/mL (Fig. 3A; 64 µg/mL: 50.1% *p*=0.0260; 128 µg/mL: 23.2% *p*=0.0072; 256 µg/mL: 12.2% *p*=0.0003). Capsaicin had no significant effect until 256 µg/mL (56.8% *p*=0.0154), with 512 µg/mL reducing transconjugants to 14.5% *p*=0.0001) (Fig. 3B). However, at high concentrations capsaicin was also noted to reduce the overall number of fluorescent cells recorded in the population, indicating the high concentrations are interfering with the fluorescence (Fig. S1).

**Figure 3.**
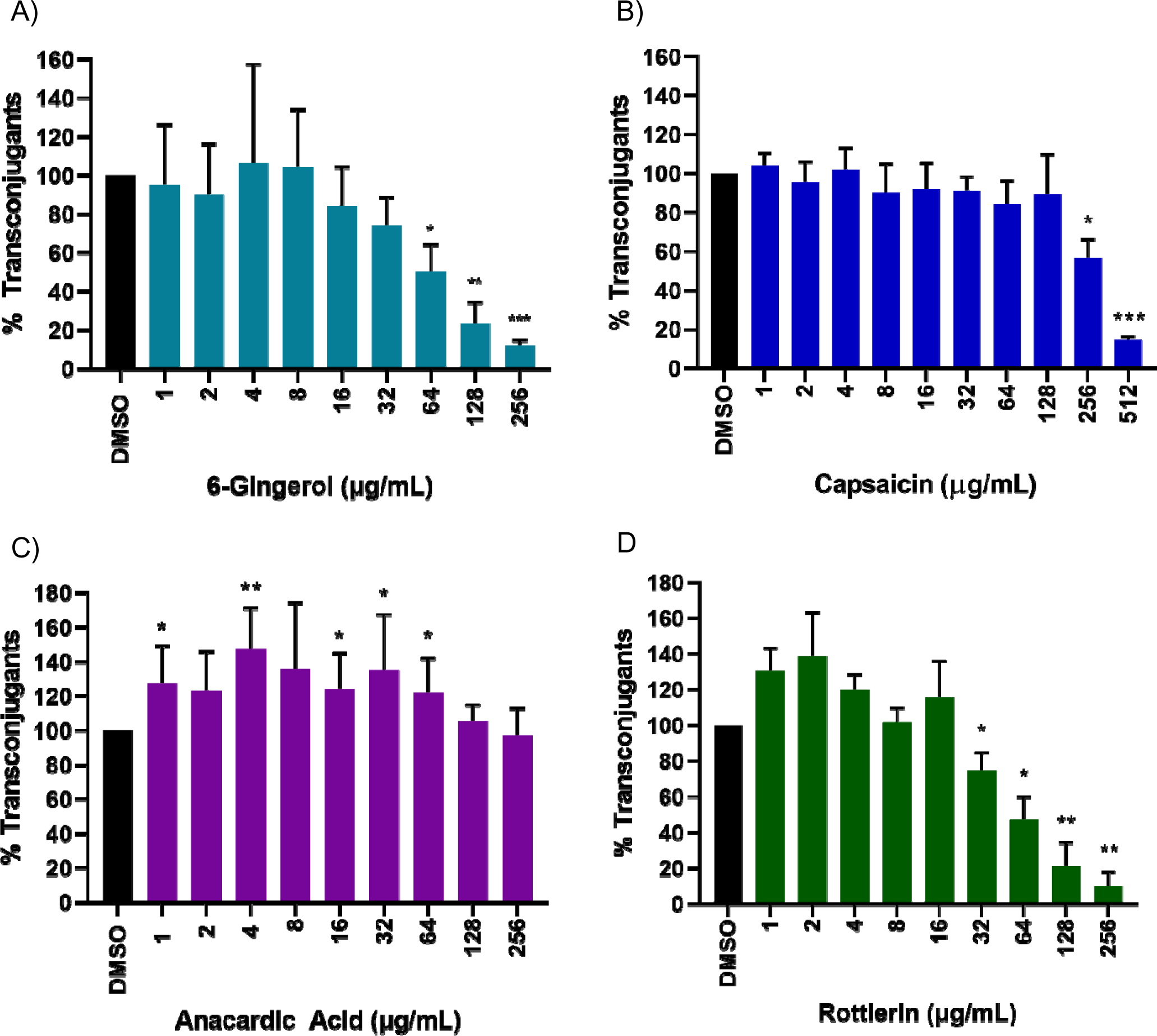
Dose-response curves of **A)** 6-gingerol on pCT transmission in *E. coli*, **B)** capsaicin on pCT transmission in *E. coli*, **C)** anacardic acid on pKpQIL transmission in *K. pneumoniae*, and **D)** rottlerin on pKpQIL transmission in *K. pneumoniae*. Data shown are the mean ± standard deviation from three independent experiments, each carried out with four biological replicates. The percentage of transconjugants following incubation with compounds was compared to a DMSO control using a paired *t*-test. Significantly different results are indicated with * (*p* ≤ 0.05), ** (*p* ≤ 0.005), or *** (*p* ≤ 0.001).

For *K. pneumoniae* Ecl8 carrying pKpQIL, two compounds were selected, rottlerin because it caused a significant decrease in transconjugant production, and anacardic acid because of the increase in transconjugant production. The results for anacardic acid were consistent with all concentrations showing an increased trend in transconjugant generation, and five of the nine concentrations were statistically significant (Fig. 3C; 1 µg/mL: 128%, *p*=0.0275; 4 µg/mL: 147%, *p*=0.0046; 16 µg/mL: 124%, *p*=0.0401; 32 µg/mL: 135%, *p*=0.0440; 64 µg/mL: 122%, *p*=0.0431). Rottlerin concentrations above 32 µg/mL reduced pKpQIL transconjugant production (Fig. 3D; 32 µg/mL *p*=0.0460, 64 µg/mL *p*=0.0182, 128 µg/mL *p*=0.0094, 256 µg/mL *p*=0.0025).

It is possible that potential anti-plasmid compounds could be inhibiting the growth of the donor or recipient strains, which would alter either the number of cells able to donate or receive the plasmid. To ascertain if this was the case with the pure compounds, antimicrobial susceptibility testing was carried out for 6-gingerol, capsaicin, anacardic acid, and rottlerin. None of the compounds inhibited the growth of the *E. coli* and *K. pneumoniae* strains (>512 μg/mL) at concentrations tested in the conjugation assays (Table 1).

**Table 1.** Susceptibility of bacterial strains to natural product compounds.

| Strain | MIC (µg/mL) |  |  |  |
| --- | --- | --- | --- | --- |
|  | 6-gingerol | Capsaicin | Anacardic acid | Rottlerin |
| *E. coli* EC958c | >512 | >512 | ND | ND |
| *E. coli* EC958c *mCherry* | >512 | >512 | ND | ND |
| *E. coli* EC958c with pCT*gfp* | >512 | >512 | ND | ND |
| *K. pneumoniae* Ecl8 | ND | >512 | >512 | >512 |
| *K. pneumoniae* Ecl8 *mCherry* | ND | >512 | >512 | >512 |
| *K. pneumoniae* Ecl8 with pKpQIL *gfp* | ND | >512 | >512 | >512 |
| *K. pneumoniae* with pCPE16_3 (clinical isolate) | >512 | >512 | >512 | >512 |
| *K. pneumoniae* ATCC 43816R (hygromycin resistant recipient) | >512 | >512 | >512 | >512 |

All strains were tested as three biological replicates. ND, not determined

### Compounds impact plasmid transmission in a carbapenem resistant *Klebsiella pneumoniae* clinical isolate

Next, the effect of the four compounds was tested on a clinical urine isolate of *K. pneumoniae*, which we have shown carries and readily transmits a 120 kb IncF plasmid with a *bla*_NDM-1_ carbapenem resistance gene, termed pCPE16_3 (19). Each compound was tested at 100 µg/mL, with a one-hour co-incubation of donors and recipients. Three parameters were explored: the absolute number of transconjugants produced at the end of the one-hour incubation, the donor-to-recipient ratio at one hour, and the conjugation frequency calculated as the number of transconjugants generated per donor after one hour. For capsaicin, there was no change in the number of transconjugant bacteria, donor-to-recipient ratio, or conjugation frequency (Fig. 4A; *p*=0.4713, 0.2513, and 0.4446, respectively). This contrasts with what was seen with pKpQIL, where the number of transconjugant bacteria increased with capsaicin (Fig. 2B). Anacardic acid did not affect the number of transconjugant bacteria, donor-to-recipient ratio, and conjugation frequency (Fig. 4B; *p*=0.0862, 0.2302, 0.1937, respectively). This was also in contrast to the results for pKpQIL (Fig. 2B). Comparable to pKpQIL, 6-gingerol had no effect on any of the parameters for pCPE16_3 conjugation (Fig. 4C; *p*=0.3988, 0.6255, 0.6850, respectively). Interestingly, rottlerin significantly reduced the total number of transconjugant bacteria from 1.32×10^5^ in DMSO to 2.25×10^4^ (*p*=0.0427), and the donor-to-recipient ratio decreased from 0.0169 to 0.00783 (*p*=0.0339) (Fig. 4D). However, the conjugation frequency did not change significantly (*p*=0.2788), likely resulting from the change in the number of donors during the experiment. To check if any of the compounds were inhibiting growth of the clinical isolate KP10 or the hygromycin-resistant recipient strain KP20, the minimum inhibitory concentration of each compound was determined (Table 1). The growth of the strains was not inhibited by any of the compounds (>512 µg/mL).

**Figure 4.**
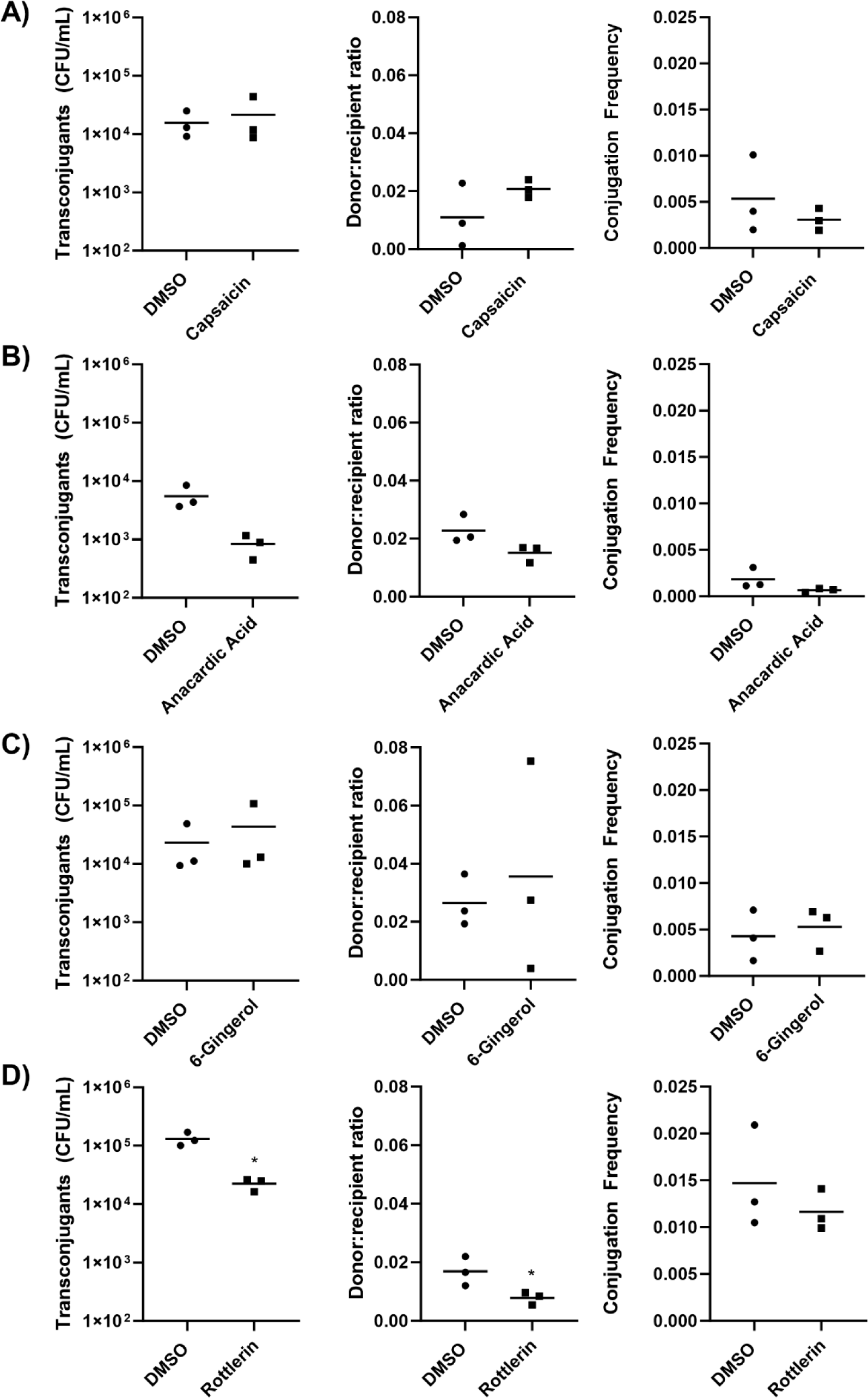
Liquid mating of the clinical *K. pneumoniae* isolate carrying the pCPE16_3 plasmid and *K. pneumoniae* ATCC 43816 (hygromycin resistant) recipient strain in the presence of pure compounds. Mixed populations were mixed with 100 µg/mL of either **A)** capsaicin, **B)** anacardic acid, **C)** 6-gingerol, or **D)** rottlerin, then incubated for one hour. For each compound, the number of transconjugants (CFU/mL), donor:recipient ratio, and conjugation frequency (transconjugants per donor) was calculated after one hour of incubation. Data shown are mean from three independent experiments, each carried out with four biological replicates. Significant differences were tested using a paired *t*-test and are indicated with * (*p* ≤ 0.05).

## Discussion

The growing threat of AMR necessitates the search for alternative strategies to combat the spread and prevalence of AMR genes. Anti-plasmid compounds that interfere with plasmid transfer or stability are being explored as a potential way to address AMR. Since natural products possess a wide range of biological activities, they offer a promising source of anti-plasmid compounds. To that end, we investigated the effect of natural product extracts and some of their reported bioactive constituents on plasmid transmission.

We found the extracts to be effective in reducing plasmid transfer in both *E. coli* and *K. pneumoniae*. To identify the possible compounds responsible for reducing plasmid transfer within the natural product extracts, we tested the major and widely reported bioactive compounds present in these extracts. We found that 6-gingerol and capsaicin were effective in reducing pCT transmission in *E. coli*. Previous work showed that 6-gingerol reduced the transfer of pKM101, pUB307, and TP114 plasmids, and capsaicin reduced the transfer of R7K, pUB307 and pKM101 plasmids in *E. coli*, without antibacterial effects on Gram-negative bacteria (20). This indicates that 6-gingerol and capsaicin may have broad-range activity against plasmid transmission in *E. coli*. Rottlerin previously reduced the transfer of pKM101, TP114, pUB307 and R6K plasmids in *E. coli* (21). Interestingly in our study, which used a clinical *E. coli* isolate with a veterinary plasmid (22), rottlerin had a minor effect on plasmid transmission, while having a strong effect in *K. pneumoniae*. In our experience, conjugation in *K. pneumoniae* is generally far less prone to interference by anti-plasmid compounds (23). This is possibly due to the presence of the thick capsule, a polysaccharide matrix, that encapsulates *K. pneumoniae* cells, making it harder for compounds to get inside cells (24). Therefore, the activity of rottlerin in this system is of significant interest and warrants further study to determine mechanism of action and full range of activity.

In our work, we used a fluorescent reporter assay to monitor plasmid transmission. Using fluorescent reporters to detect transmission events increases throughput, however, consideration must be given to account for the potential impact of compounds on the expression and function of fluorescent proteins. For example, with coloured extracts and compound treatment, some of which are bright red/orange, a change in the baseline fluorescence of cells can occur. This reduction in fluorescence corresponds to the apparent reduction in the number of transconjugants observed. This impact on fluorescence may also account for the apparent discrepancy between the screening of capsaicin at 100 μg/mL and the dose response curve, where activity was not detected until 256 μg/mL.

The activity of anti-plasmid compounds could also be influenced by plasmid-host combination. Some bacterial host strains can acquire and maintain certain plasmids without a fitness cost (25). These successful host-plasmid pairings significantly contribute to the global spread of AMR genes (26). Despite their prevalence, there is still limited knowledge on what factors influence plasmid transfer in these successful plasmid-host combinations (27). Therefore, understanding the intricacies of the plasmid-host relationship is important in developing effective strategies to combat plasmid-mediated antibiotic resistance. The ideal anti-plasmid compound would have broad-range activity against different host strains and plasmids, however, owing to the diversity of plasmids and their relationship with the host, this may prove difficult. Nonetheless, identifying compounds that target globally disseminating plasmid-host combinations could be an attractive strategy to prevent the spread of AMR plasmids.

The natural product extracts showed very strong activity against transmission of both plasmid/host strains, while the pure compounds tested had different levels of activity. This indicates there are still unidentified compounds present in the complex extracts. Further work could test fractions of the extracts and/or additional pure compounds for anti-plasmid activity, as the pure compounds we have tested do not account for the entirety of the activity seen in extracts. It is also possible that distinct combinations of compounds are responsible for the strong anti-plasmid activity seen in the extracts.

## Methods and materials

### Natural product extracts and compounds

Extracts were produced by extracting 10 g of powdered plant material with chloroform (200 mL) overnight at room temperature. Extracts were then concentrated under vacuum and stored in a freezer at -20 °C until use. Compounds 6-gingerol, capsaicin, anacardic acid, and rottlerin were purchased from Merck, UK. Extracts or compounds were dissolved in DMSO, and DMSO controls were prepared to the same concentration and used throughout.

### Bacterial strains

Bacterial strains used are described in Table 2. Unless stated otherwise, all strains were grown in Luria-Bertani (LB) broth and incubated at 37 °C with aeration.

**Table 2.**
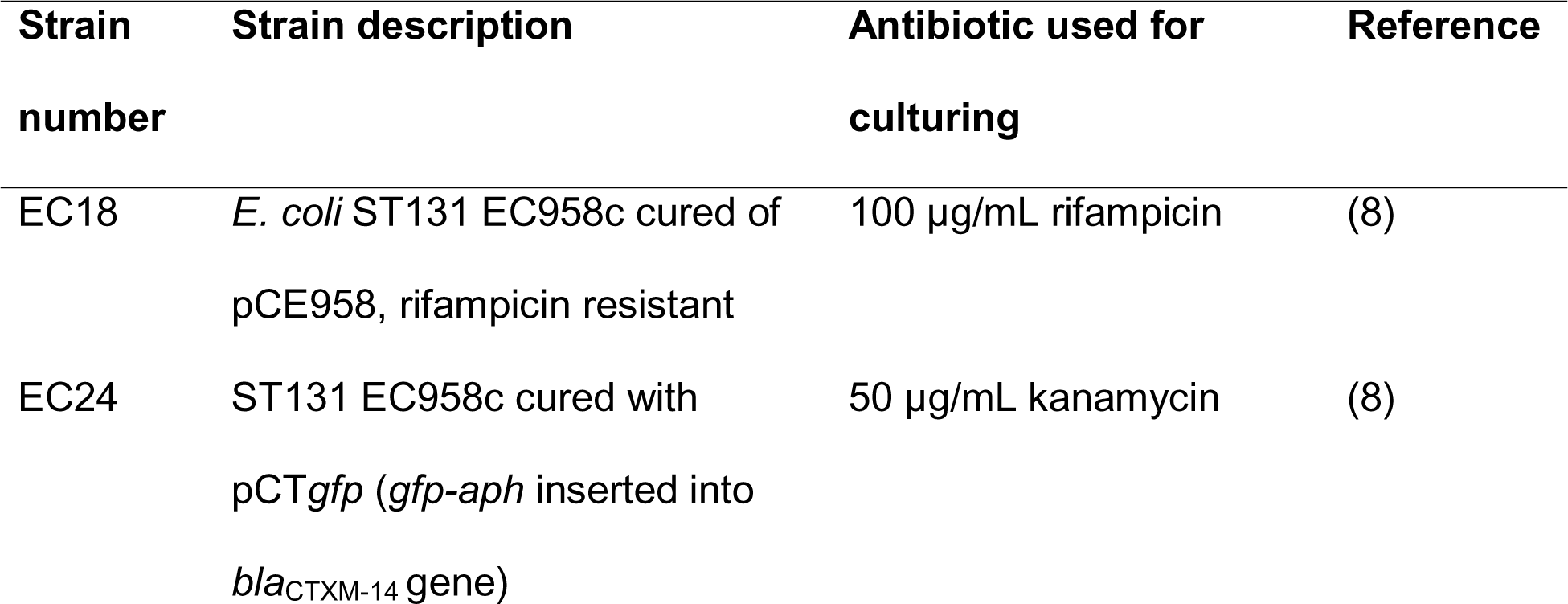
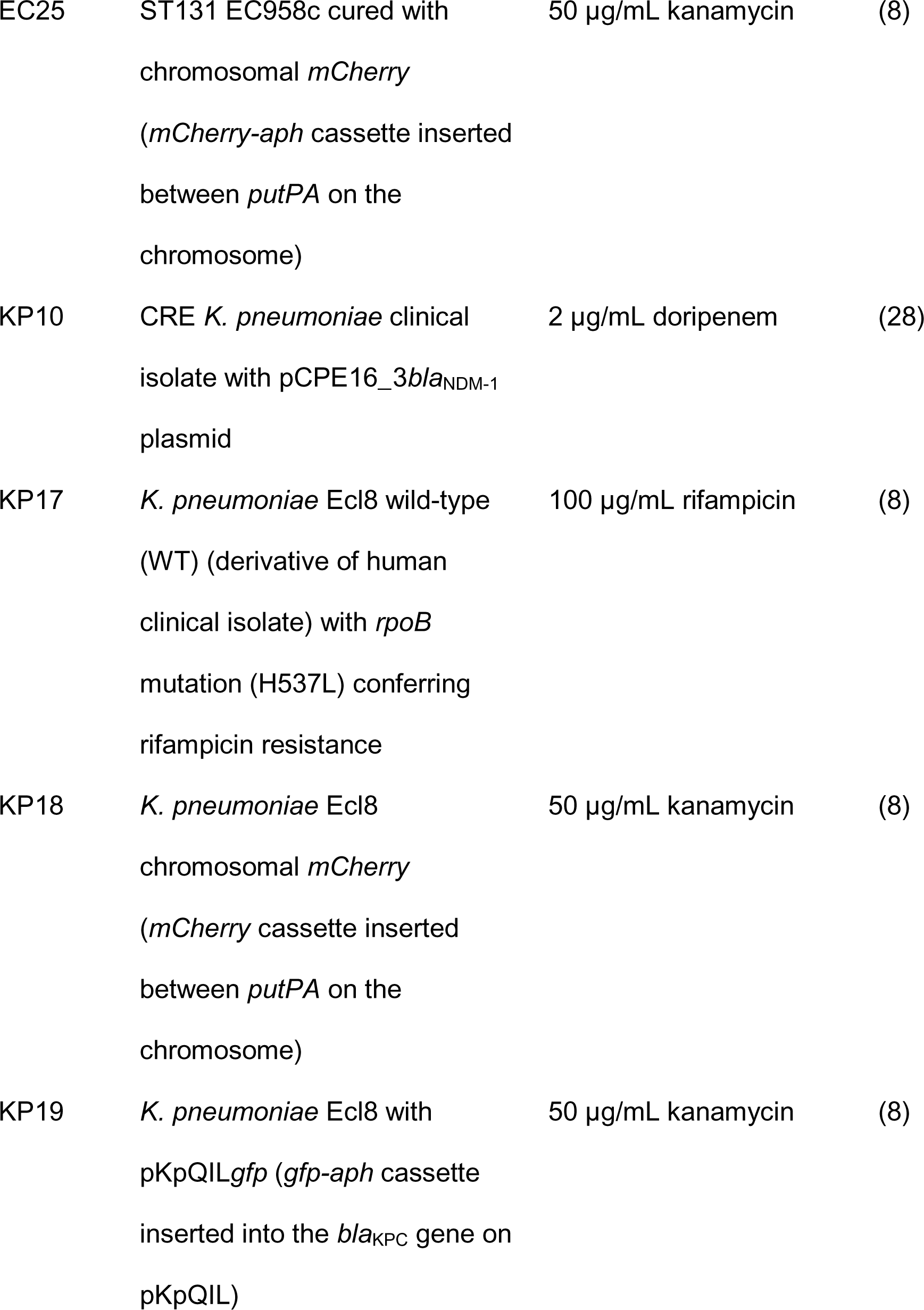
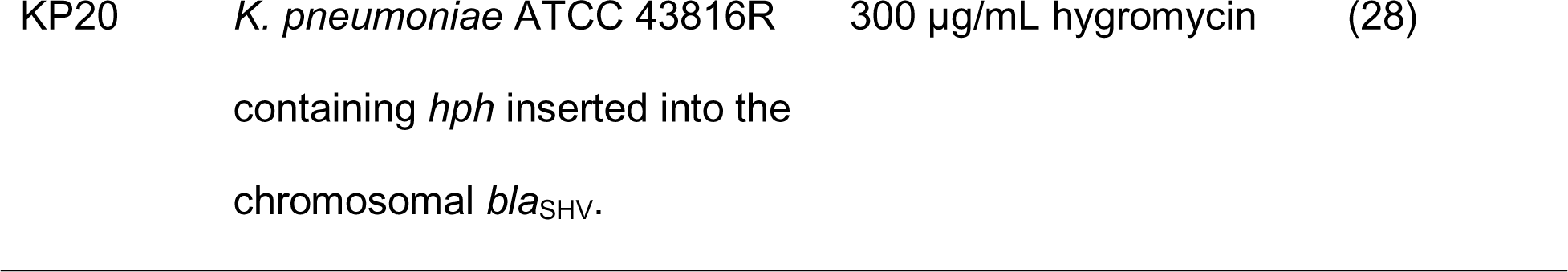
Bacterial strains and antibiotics used for routine culturing.

### High-throughput screening of extracts and compounds

The transmission of pCT*gfp* in *E. coli* ST131 EC958c and pKpQIL*gfp* in *K. pneumoniae* Ecl8 in the presence of natural product extracts and compounds was measured by flow cytometry as previously described (8), without any modifications.

### Antimicrobial susceptibility testing

The antimicrobial susceptibility of bacterial strains to compounds were determined using the broth microdilution method as previously described (29).

### Liquid broth conjugation with clinical isolate

The *K. pneumoniae* clinical isolate carrying pCPE16_3*bla*_NDM-1_ plasmid (KP10) was paired with the hygromycin resistant *K. pneumoniae* ATCC 43816R recipient strain (KP20) in liquid broth as previously described (28). Briefly, KP10 and KP20 cultures were grown overnight, sub-cultures were prepared in 5 mL LB broth (1% inoculum) and grown to an OD_600_ of ∼0.5. After, 1 mL of cultures was pelleted, and media were replaced with LB broth to adjust the OD_600_ to 0.5. The donor KP10 and the recipient KP20 were mixed at a 1:10 ratio alongside control single strains. The donor (KP10), recipient (KP20), and mixed cultures were separately diluted 1:5 in LB broth containing a final concentration of 100 µg/mL of the natural compound or 100 µg/mL DMSO as vehicle control and these were incubated statically at 37 °C for one-hour. Corresponding dilutions were plated onto LB agar to assess cell viability and selective media to determine donor-to-recipient ratios and transconjugant production. Plates were incubated at 37 °C overnight. Transconjugant colonies carrying pCPE16_3*bla*_NDM-1_ were selected on LB agar supplemented with 300 µg/mL hygromycin B (PhytoTech Labs, USA) and 2 µg/mL doripenem (Merck, Germany). Conjugation frequencies were calculated using the following formula:

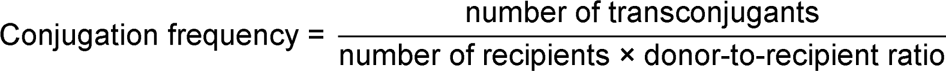

Data shown are the mean ± standard deviation of three independent experiments, each carried out with four biological replicates.

